# Intrinsic linking of chromatin fiber in human cells

**DOI:** 10.1101/2022.07.13.499767

**Authors:** Maciej Borodzik, Michał Denkiewicz, Krzysztof Spaliński, Kamila Winnicka-Sztachelska, Kaustav Sengupta, Marcin Pilipczuk, Michał Pilipczuk, Yijun Ruan, Dariusz Plewczynski

**Affiliations:** Institute of Mathematics, University of Warsaw, ul. Banacha 2, 02-097 Warsaw, Poland; Centre of New Technologies, University of Warsaw, ul. Banacha 2c, 02-097 Warsaw, Poland; Faculty of Mathematics and Information Science, Warsaw University of Technology, Warsaw, Poland; Institute of Informatics, University of Warsaw, ul. Banacha 2, 02-097 Warsaw, Poland; The Jackson Laboratory for Genomic Medicine, USA

## Abstract

**Motivation:** We propose a practical algorithm based on graph theory, with the purpose of identifying CTCF-mediated chromatin loops that are linked in 3D space. Our method is based finding clique minors in graphs constructed from pairwise chromatin interaction data obtained from the ChIA-PET experiments. We show that such a graph structure, representing a particular arrangement of loops, mathematically necessitates linking, if co-occurring in an individual cell. The presence of these linked structures can advance our understanding of the principles of spatial organization of the genome.

**Results:** We apply our method to graphs created from in situ ChIA-PET data for GM12878, H1ESC, HFFC6 and WTC11 cell lines, and from long-read ChIA-PET data. We look at these datasets as divided into CCDs - closely interconnected regions defined based on CTCF loops. We find numerous candidate regions with minors, indicating the presence of links. The graph-theoretic characteristics of these linked regions, including betweenness and closeness centrality, differ from regions without, in which no minors were found, which supports their non-random nature. We also look at the position of the linked regions with respect to chromatin compartments.

**Availability:** The implementation of the algorithm is available at https://github.com/SFGLab/cKNOTs

## 1 Introduction

The human genome is composed of more than 3 billion nucleotides, measures about 2 meters and it is enclosed in the nucleus that is only 6 micrometers wide (Tang, et al., 2015), causing the spatial structure of the genome to emerge. This spatial organization includes multiple levels of interrelated structures (Wang, et al., 2016), most importantly: chromatin compartments, topologically associated domains or TADs (Dixon, Gorkin, & Ren, 2016), and chromatin contact domains or CCDs (Tang, et al., 2015); for an illustration see Figure 1A.

**Figure 1:**
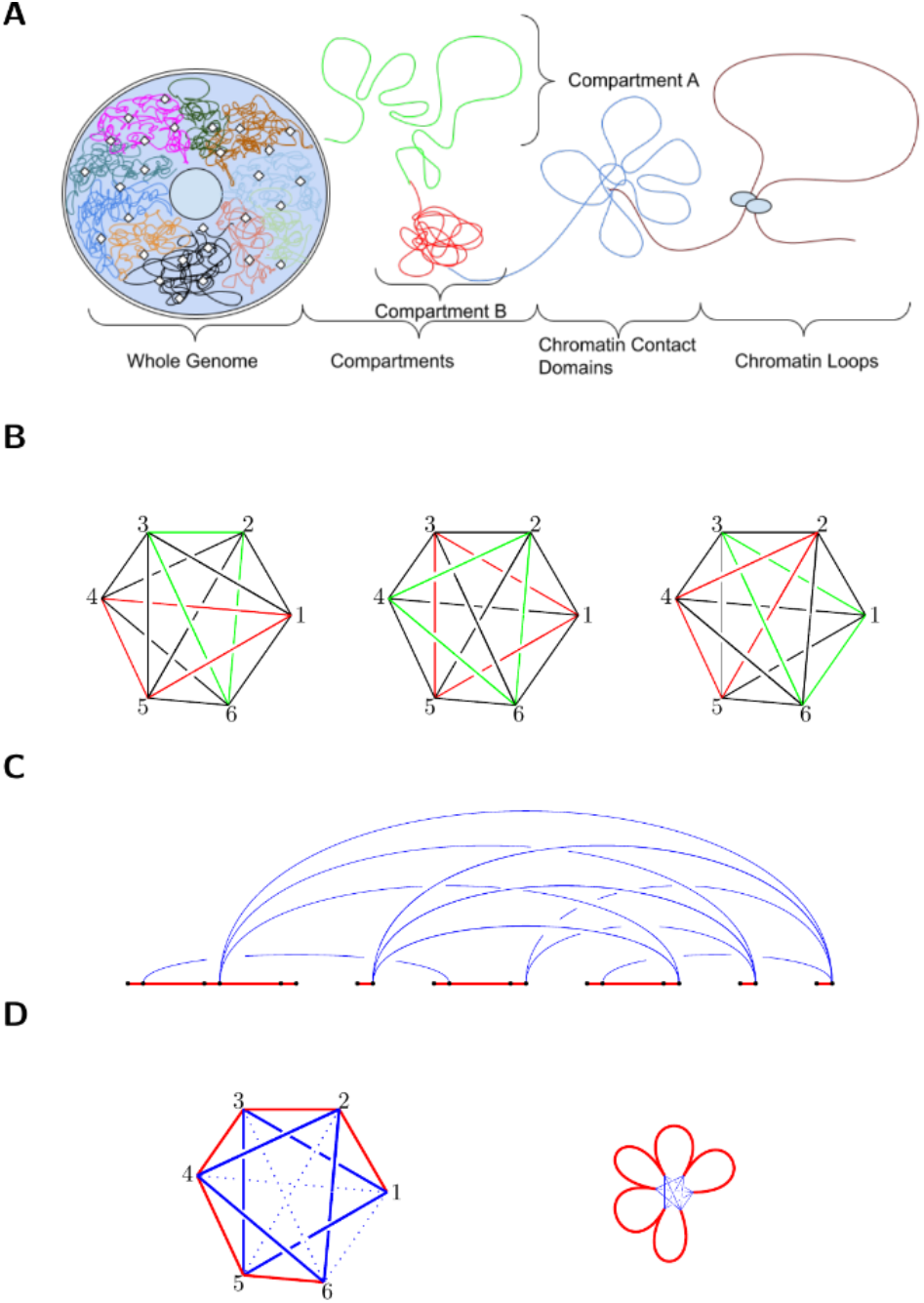
A) Illustration of the hierarchical structure of the genome. B) Different geometric realizations of K_6_ graph lead to different pairs of triangles that are linked. Observe that the central picture and the right one differ in a single detail, the line connecting 2 and 5 is above or below the line connecting 1 and 5, but the pairs of triangles are different. C) An example of a linear minor. The blue arcs represent ChIA-PET contacts (jump edges), the red lines and dots denote the node sets being collapsed to a minor. D) Illustration of the necessity of using only solid minors. The linking of a blue solid triangle (formed of jump edges) has no biological meaning. After contracting blue edges to small size, there is no linking in the DNA thread. Dotted lines represent other edges, that are not the focus here

The properties of the genome structure (e.g. boundaries between TADs) are partially determined by the formation of chromatin loops, which can be detected using chromatin interaction analysis with paired-end tags (ChIA-PET) experiment (Li, et al., 2010). Such loops comprise the Cohesin protein complex bringing together in 3D space two occupied CTCF (CCCTC-binding factor) binding sites, which can be far apart in terms of genomic coordinates, i.e., the linear position on the DNA strand (Splinter, et al., 2006) (Ong & Corces, 2014). They are also used to define CCDs as continuous regions with a relatively large number of interconnected loops (Tang, et al., 2015). The 3D structure changes resulting from the formation or disruption of the loops were found to have functional consequences (Li, et al., 2012) (Doyle, Fudenberg, Imakaev, & Mirny, 2014) (Dixon, Gorkin, & Ren, 2016).

Another important level of segmentation, which controls the functionality in the genome, is compartmentalization. The genome is divided into two major compartments: A and B. Compartment A, or *euchromatin*, is open for transcription to take place, and therefore active. On the other hand, compartment B, called *heterochromatin*, is densely packed, so that transcription is suppressed as transcription factors can’t reach the genes. The two compartments can be further subdivided based on their location. Within compartment A we have regions denoted as A1, which are close to nuclear speckles, and A2, which is enriched with Pol II and Bromodomain Proteins and far from the nuclear speckles. Similarly, compartment B is divided into three sub-compartments: B1 is binded by Polycomb repressive complex (PRC), B2 is near the nucleus and bounded by nuclear-associating domains (NADs) and heterochromatin proteins (HP1), and B3 is close to nuclear lamina and bounded with Lamina-associating domains (LADs) along with HP1.

We know that the organization of the genome plays a significant role in its function, and disturbances in chromatin folding influence gene expression (Lupiáñez, et al., 2015). The principle of creating this organizational complexity is not fully understood (Szalaj & Plewczynski, 2018). In particular, one unexplored possibility is that chromatin loops might be linked together, impacting the resulting 3D structure. Current methods of analyzing chromatin conformation, such as Chromatin Interaction Analysis by Paired-End Tag Sequencing (ChIA-PET), (Fullwood, et al., 2009) or Hi-C (Lieberman-Aiden, et al., 2009), give us point-wise information about which pairs of loci are closely placed in 3D space. Nevertheless, there are many possible 3-dimensional models corresponding to this data, therefore it is still complicated to decide which one of them is accurate. Graph theory can aid in solving this problem by abstracting from the particularities of any 3D physical model and looking only at connection structure. A continuous region of chromatin (a chromosome or its part) can be represented as a graph composed of vertices, which represent points on the genome, and edges, which correspond to either to interactions (physical connection between distant loci) detected using conformation capture experiments, or to the parts of DNA strand itself.

Although we do not know which spatial structure of chromatin is correct, in the graph representation we can find features common for every possible model compatible with data.

We describe a computational method of searching for so-called *minors* of a complete subgraph with 6 vertices (K_6_) within the chromatin graphs. The presence of such a minor, guarantees that the chromatin graph embedded in 3-dimensional space, contains two linked loops. The base (exact) algorithm has exponential complexity; hence we refine it to only search for a certain subset of K_6_ minors, which we call linear minors. We apply the linear algorithm to graphs constructed from ChIA-PET data for GM12878 and three other cell lines, subdivided into densely connected regions called chromatin contact domains, or CCDs (Tang, et al., 2015). After identifying the linked regions, we study the graph-theoretic properties of the corresponding graphs, to learn their characteristics. We also provide a 3D model of an example region on chromosome 10, containing a K_6_ minor. Finally, we assess the coverage of each CCD-containing minor by the two major chromatin compartments.

## 2 Methods

### 3.1 Datasets

To create the graphs representing the physical interactions in chromatin, we used data from several ChIA-PET experiments targeting the CTCF protein. We use data from two versions of ChIA-PET protocol: the long-read ChIA-PET (Li, et al., 2017), and the in situ ChIA-PET: a more efficient version detecting interactions directly in the nucleus.

We applied our algorithm to in situ CTCF ChIA-PET datasets for 4 human cell lines: GM12878 (human lymphoblastoid cell), H1ESC (H1 human embryonic stem cell), HFFC6 (human foreskin fibroblast cell) and WTC11 (human induced pluripotent stem cell), and to long-read CTCF ChIA-PET for GM12878 cell line. The in situ ChIA-PET data was obtained from 4DNucleome data portal (Reiff, et al., 2021) (Dekker, et al., 2017) and has been produced by the Jackson Laboratory. The long-read data was provided by (Tang, et al., 2015). The in situ ChIA-PET data is mapped onto hg38 reference genome, while the long-read data was mapped to hg19 reference genome. The raw interactions in ChIA-PET data are provided as pairs of anchors: each having a start and end coordinate, and the frequency of occurrence of the given interaction, called the PET-count. In other words, the presence of an interaction between two loci in this dataset means that at least two reads existed, whose both end-points were approximately the same, and the endpoints were confirmed to be CTCF binding sites - verified by both the presence of a peak in the ChIP-Seq signal (indicating the presence of CTCF protein) and the presence of a CTCF binding motif in the DNA sequence. The raw interactions were filtered by their frequency (PET-count), and only the ones with frequency equal or above a threshold were retained. The threshold was set to 3 for in situ data (for all cell lines), and 2 for long-read data - this is because the latter dataset is generally sparser. Note, that the ChIA-PET read frequency should not be understood as an absolute estimate of probability of the existence of a loop in an arbitrary cell. The detection of an existing chromatin loop via ChIA-PET read is itself highly chance-dependent, while the PET-counts themselves depend on e.g., sequencing depth. Thus, only cautious relative inferences can be made, with higher PET-count interactions being regarded as more confident than those with lower PET-count. Next, we merged all overlapping anchors, as we are interested in the case of multiple loops attached to (almost) the same anchor region.

At this stage we could proceed with graph construction, producing a single connected graph for each chromosome. However, since it is not feasible to run the algorithm on whole-chromosome networks, we extract interactions for each individual CCD separately. Since the CCDs are the most internally connected regions by definition, it makes most sense to search for linking within them.

### 3.2 Graph construction and graph-theoretic definitions

#### Chromatin graph representation

We represent the interaction data of a continuous segment (such as a CCD) of the human genome as a connected graph G, with vertices V and edges E. The vertices V = v_1_, v_2_, …, v_n_ are placed on the chromatin strand in this order, each representing a genomic locus. The edges belong to one of two classes: *solid edges* or *jump edges*. The edges E_s_ = v_i_v_i+1_ for 1 ≤ i < n connect consecutive vertices and represent the DNA strand. The remaining set of the edges E_j_ we will call *jump edges*. A *jump edge* corresponds to the presence of a chromatin loop joining two distant points of the chromatin strand. In our model, such interaction is reflected by a single jump edge, connecting the corresponding vertices v_j_ and v_k_. The actual coordinates which we choose to represent are determined by the endpoints of jump edges. The jump edges are quite short in terms of genomic coordinates in relation to the chromosome scale, as most of them connect parts of genome up to 1 million base pairs away. However, in 3D space they bring the two endpoints to close proximity. Within graphs constructed in this way we search for linked structures, which we will now define.

#### Graph minors and linking in a graph

A *geometric realization* of a graph G is a particular arrangements of its vertices and edges in 3D space. Formally, it is an assignment of each vertex v ∈ V(G) a point φ(v) in R^3^, and to each edge uv ∈ E(G) an arc in R^3^ with endpoints φ(u) and φ(v). We require that no two arcs intersect, except possibly at their common endpoints. A geometric realization is linked if there are two cycles in C_1_ and C_2_ in G, whose combined realizations φ(C_1_) ∪ φ(C_2_) form a non-split link in S^3^. Intuitively, it means that the curves in 3D space that are the cycles’ representations are interleaved with one another so that they cannot be pulled apart.

Given a geometric realization, we say that the graph G is *linked*, if there exist two cycles in G, whose geometric representations in this realization are linked. Finally, if a graph is linked for any possible of its geometric realizations, it is called and *intrinsically linked graph*. An important observation here is that K_6_ (i.e. the clique graph on six vertices) is intrinsically linked. To better understand the concept of linkedness, we provide the example of different geometric realizations of the K_6_ graph depicted in Figure 1B.

Intrinsically linked graphs were characterized by Conway and Gordon (Conway & Gordon, 1983) in terms of minor containment, which we will now briefly recall. Given graphs H and G, a *minor model* of H in G is an assignment to every v ∈ V(H) a connected subgraph F_v_ of G such that (1) the subgraphs F_v_ are pairwise vertex disjoint, and (2) for every uv ∈ E(H) there is at least one edge in G between V(F_u_) and V(F_v_). Minors can be intuitively understood in the following way: G has a minor model of H if we can obtain H from G by contracting edges (i.e., for and edge *uv* replacing *u* and *v* with a single vertex which retains the neighbors of *u* and *v*) and possibly removing some vertices and edges. Here we will only consider *clique minors* where H is a clique K_r_ on r vertices. We provide a general description for any r, but in our empirical work we use r = 6, in other words we consider K_6_ minors. Note that for the sake of conciseness we will often use the word “minor” to refer to a minor model.

A graph can have many different K_6_ minor models. In that case any of its geometric realizations could have multiple pairs of linked cycles. If two K_6_ minor models in a graph partially overlap, the corresponding pairs of linked cycles might, but need not, be the same.

The central theorem related to our method, formulated by Conway, states the following: A graph G is intrinsically linked if and only if it contains a minor model of K_6_. We can thus detect linked structures by finding K_6_ minor models in the respective chromatin graphs. Given the importance of minors in our study, we introduce a notation for them: as alluded to above, a K_6_ minor consists of connected subgraphs H_1_,…,H_6_ which are connected and pairwise disjoint, as well as the choice of edges e_ij,_ 1 ≤ i < j ≤ 6, such that e_ij_ connects a vertex in H_i_ with a vertex in H_j_.

#### Solid minors

There is one more thing to consider: not every K_6_ minor model in such a graph will lead to the linking of the chromatin strand. The problem is that linking cycles consisting only of jump edges would not result in a biologically meaningful linked structure - the cycles can be linked but there is no linking of the actual physical chromatin molecule. A schematic of such a situation is presented in Figure 1D. To address this we are looking for a certain type of K_6_ minor models, which we will call *solid minors*. Let G be a graph representation of chromatin as defined above. A cycle in G is *solid* if it contains at least one edge of E_s_ (i.e., part of the chromatin strand).

Recall, that since K_6_ is intrinsically linked, for any geometric realization of K_6_ graph with the vertex set V(K_6_) = {1, 2, 3, 4, 5, 6} there exists a triple of vertices a, b, c, such that the triangle abc and the triangle formed by the three other vertices have linked geometric realizations. Let M = (F_v_) for v ∈ V(K_6_) be a K_6_-minor model in G. Then M is *solid* if for every triangle uvw of K_6_ there exists a solid cycle in G that traverses F_u_, F_v_, F_w_, and exactly one edge between each pair of these subgraphs. To summarize any solid minor of a graph G leads to an actual linking of the chromatin strand, and thus detecting solid minors is the focus of our method.

### 3.3 Treewidth, pathwidth, and cutwidth

Before describing the algorithm for finding solid minors we will give a few more definitions, of which the most important in this work is *path decomposition* and *pathwidth*, which can be introduced as simple special cases of *tree decomposition* and *treewidth* respectively. Treewidth, (Robertson & Seymour, 1984) is arguably one of the most successful structural graph parameter. Low treewidth means that a graph structurally resembles a tree. Coincidentally a vast number of fundamental hard computational problems becomes tractable on graphs of low treewidth; we refer to Chapter 7 of (Cygan, et al., 2015) for an overview. Algorithms on graphs of bounded treewidth usually follow the paradigm of dynamic programming and, consequently, are very robust with regards to different variants of the studied problems. The problem of finding minor models is no exception.

Let us proceed with formal definitions. A *tree decomposition* of a graph G consists of a tree T and a function β : V(T) → 2^V(G)^ that assigns to every node t ∈ V(T) a *bag* β ⊆ V(G). The bags are required to satisfy the following two properties: (i) for every v ∈ V(G) the set of nodes {t ∈ V(T) | v ∈ β(t)} induces a nonempty connected subtree of T, and (ii) for every edge uv ∈ V(G) there exists a node t ∈ V(T) with {u, v} ⊆ β(t). The width of the decomposition (T, β) equals max_t ∈ V(T)_ |β(t)| - 1 and the treewidth of a graph is the minimum possible width of its tree decomposition.

The crux of the definition of a tree decomposition is that for every edge st ∈ E(T) the set β(s) ∩ β(t) is a separator between the vertices of G appearing in the bags of the two connected components of T-{st}. This allows dynamic programming algorithms that scan the tree T in a bottom-to-top fashion.

If one restricts the above definition to T being a path (as opposed to an arbitrary tree), one obtains the definition of a *path decomposition* and *pathwidth*. While the value of pathwidth can be larger than that of treewidth, dynamic programming on path decompositions is often conceptually and technically simpler than on tree decompositions. Path decomposition is directly applied in our algorithm.

Finally, *Cutwidth* is another graph width parameter that is relevant to our work. Let G be a graph and let ≺ be a total order on V(G). For every v ∈ V(G), the *cut* at v on ≺ is defined as the partition V(G) = A_v_ ∪ B_v_ where A_v_ = {u ∈ V(G) | u ≼ v} and B_v_ = {u ∈ V(G) | v ≺ u}. The *order* of the cut A_v_, B_v_ is the number of edges of G with one endpoint in A_v_ and the second endpoint in B_v_. The *cutwidth* of the ordering ≺ is the maximum order of the cut on ≺ and the cutwidth of a graph is the minimum cutwidth among all orderings of V(G).

The starting point of our work is an observation that in a chromatin graph G with V(G) = {v_1_, v_2_, …, v_n_} with the total order v_1_ ≺ v_2_ ≺ … ≺ v_n_ (which is the natural ordering of vertices by their actual genomic coordinates of each vertex) is likely to have low cutwidth. This is because of the characteristics of chromatin organization, especially the existence of contact domains – since most edges (interactions) are restricted to these domains, the number of edges stacked in one location is limited. This observation is supported by the experimental results, for example in long-read ChIA-PET for GM12878 data, the cutwidth ranges between 15 (chromosome 21 and 22) and 31 (chromosome X).

However, the values of cutwidth are still quite large for dynamic programming algorithms whose running times depend exponentially on the value of the width parameter. Therefore, we study as well the pathwidth of the chromatin graphs. To this end, we need the following well-known observation. Let G be a graph and let ≺ be a total order of V(G) of cutwidth *c*. Consider the following path decomposition (T, β) of G. Assume that V(G) = v_1_, v_2_, …, v_n_ with v_1_ ≺ v_2_ ≺ … ≺ v_n_. Let T be a path with vertices labeled 1, 2, …, n in this order and let β(i) = {v_i_} ∪ {j < I | ∃_k ≥ i_ v_j_v_k_ ∈ E(G)}. It is straightforward to verify that (T, β) is a path decomposition of width at most *c*.

Using the observation above we turned the natural ordering v_1_ ≺ v_2_ ≺ … ≺ v_n_ of a chromatin graph into a path decomposition. The example calculation results on the same dataset show pathwidth between 9 (chromosome 18 and 21) and 24 (chromosome X, but next largest is 16 for chromosome 13). The pathwidth bounds are much lower than the cutwidth bounds (difference ranges from 5 up to 19), indicating that often many edges in cuts 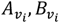 have common endpoints. These path decompositions are used as inputs for the implemented dynamic programming algorithms.

### 3.4 Finding minors via dynamic programming

The base algorithm is a straightforward procedure that tries to construct a minor model of a clique K_r_, given a path decomposition of a graph G. Let us denote V(K_r_) = {w_1_,w_2_, …, w_r_}. Given a bag β(t), a single *state* consists of

1. a function f: β(t) → {0, 1, 2, …, r},
2. a partition p_i_ of f^-1^(i), for each i ∈ {0, 1, 2, …, r}, and
3. a subset 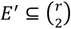

In simple terms, the state stores a 2-way map between each vertex from the bag β(t) and a vertex from K_r_ (pt. 1, 2), and a subset of edges from K_r_ (pt. 3). For a state 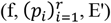, we keep a Boolean variable that indicates if one can construct a partial minor model of K_r_ from the vertices of the graph appearing in the bags to the left of the bag β(t) such that f^-1^(i) are exactly the vertices β(t) that are in 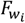 with p_i_ being the partition of them among the connected components in the partial minor model and E’ is the set of pairs 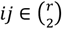 for which an edge joining 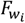 with 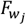 has already been found. Observe that for a given bag β(t) of size k the number of states is bounded by 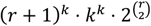, where the first factor corresponds to the number of possible functions f, the second one bounds the number of partitions 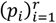 and the third one is the number of possible sets E’.

Using this algorithm we were able to find multiple K_6_ minors in the chromatin graphs. However, the above algorithm is not able to find *solid* clique minors without significant enhancement. If one does such enhancement directly, then one needs to store for every distinct u, v ∈ β(t)-f^-1^(0) the information whether one can connect u and v within the model using only vertices of 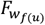 and 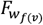 and at least one edge of E_s_. This increases the number of states to exponential in Ω(|β(t)|^2^), making the application of the algorithm infeasible in practice.

To cope with this problem, we restrict our search to a subclass of solid minor models of a clique K_r_, which we will call *linear minors*, with the property that vertices from G mapping to a single K_r_’s vertex must form continuous, non-interleaving groups on the chromatin strand.

Formally, if G is a chromatin graph with V(G) = v_1_, v_2_, … v_n_, then a *linear model* of K_r_ consists of (1) subgraphs F_i_ for i = 1, …, r where F_i_ is the subpath between 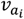 and 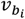 we require a_i_ ≤ b_i_ for every 1 ≤ i ≤ r and a_i+1_ = b_i+1_ for every 1 ≤ i < r, and (2) selected jump edges e_i,j_ with endpoints in F_i_ and F_j_ for every 1 ≤ i ≤ r and 1 ≤ j ≤ r with i + 2 ≤ j. A linear model is *solid* if no three edges e_i,j_ form a triangle; note that this corresponds to the minor model (in the usual sense) being solid when the edge e_i,j_ is always used to connect subgraphs F_i_ and F_j_ for the purpose of building solid cycles.

We look for linear models of K_6_. To this end, we use the aforementioned path decomposition of a chromatin graph G constructed from the ordering v_1_ ≺ v_2_ ≺ … ≺ v_n_. Recall that in this context the bag β(a) for 1 ≤ a ≤ n consists of v_a_, v_a-1_, and all vertices v_b_ for b < a that are incident with some jump edge v_b_v_c_ for some c ≥ a. A state for a bag β(a) now consists of a partial linear minor model defined as follows:

1. an index 1 ≤ ι ≤ 6 such that v_a-1_ ∈ F_ι_,
2. choice of some of the edges e_i,j_ for every 1 ≤ i ≤ ι and ι ≤ j ≤ 6 with i + 2 ≤ j if their left endpoints are to the left of v_a_.

While building the partial linear minor models in the dynamic programming algorithm we ensure that the edges e_i,j_ never form a full triangle. Note that in pt. 2 of the state for K_6_ one needs to store up to 9 edges for ι ∈ {3,4} (e.g., for ι = 3, one may need to store e_1,3_, e_1,4_, e_1,5_, e_1,6_, e_2,4_, e_2,5_, e_2,6_, e_3,5_, e_3,6_). This, if the order v_1_ ≺ v_2_ ≺ … v_n_ has cutwidth c, then we have an upper bound of 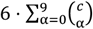 on the number of states.

### 3.5 Topological measures

In order to characterize the graph topology between the regions containing links, we quantify the size of each CCD graph in terms of vertex and edge counts, density (ratio of actual edge number and largest possible edge number). Moreover, for each node we compute two arguably most commonly used centrality measures: *closeness centrality* and *betweenness centrality* - quantities understood to indicate the importance of a given node in a graph. We will now briefly recall their definitions.

The *closeness centrality* of a node u is the inverse of the mean distance between u and all other nodes in a graph: 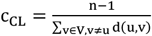, where distance d(u, v) is the length of the shortest path between u and v, and n is the number of nodes in the graph. Closeness centrality was originally defined simply as the reciprocal of the sum of the distances from u to all other nodes (Sabidussi, 1966), but we use the normalized version in which the maximum value is 1. Note, that in our case the graphs are always connected, so d(u, v) is always well-defined.

The *betweenness centrality* of a node indicates how often a node is encountered on a shortest path between other nodes, capturing the notion of being an intermediary between them (Freeman, 1977). In formal terms: 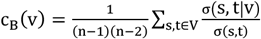 where V is the set of nodes, σ(s, t) is the number of shortest paths between s and t, and σ(s, t | v) is the number of those paths passing through node v, assuming σ(s, t) = 1 in case of s = t. In the case of multiple shortest paths, the proportion of them passing through u is considered. Again, we use the normalized version of the measure, in which values range from 0 to 1.

The centrality measures are defined per node, but their aggregates can be used to characterize the graphs themselves (Zubek, et al., 2017). In this vein we compute average values of the per-node statistics (betweenness, closeness, and the degree) thus obtaining three measurements for each CCD graph. Finally, we look at the size of CCD in terms of genomic coordinates. Using all these measures we can compare the CCDs containing and not containing minors in terms of their general topology.

## 3 Results and Discussion

For the GM128787, a total of 1091 unique solid minors were found in the long-read data, occupying 382 (16.7%) out of 2290 CCDs. In the case of in situ ChIA-PET, 725 unique solid minors were detected in 257 out of 2466 CCDs (10.2%). For other cell lines the percentages are 10.3% (HFFC6), 6.8% (WTC11) and 1.0% (H1ESC). This percentage, however, is expected to be influenced by the density of the graphs obtained from ChIA-PET data, which in turn depends on many factors, such as on the characteristics of the experiment (ling-read vs in-situ ChIA-PET), and on the PET-count threshold used. Moreover, each dataset has a distinct set of CCDs. The detailed numbers of unique solid minors located on each chromosome are provided in Table 1, while the statistics for CCDs occupied by minors are shown in Table 2.

**Table 1:** Number of unique solid linear minors in each chromosome found by the linear algorithm for each of the datasets.

|  | GM12878(LR) | GM12878(IS) | H1ESC | HFFC6 | WTC11 |
| --- | --- | --- | --- | --- | --- |
| chr1 | 71 | 36 | 24 | 44 | 123 |
| chr2 | 134 | 73 | 4 | 79 | 54 |
| chr3 | 89 | 71 | 1 | 23 | 21 |
| chr4 | 118 | 26 | 0 | 46 | 0 |
| chr5 | 101 | 52 | 4 | 105 | 1 |
| chr6 | 59 | 18 | 0 | 35 | 5 |
| chr7 | 94 | 58 | 0 | 21 | 4 |
| chr8 | 57 | 28 | 0 | 31 | 15 |
| chr9 | 51 | 42 | 0 | 20 | 5 |
| chr10 | 14 | 39 | 0 | 90 | 13 |
| chr11 | 28 | 31 | 0 | 10 | 1 |
| chr12 | 57 | 53 | 0 | 58 | 3 |
| chr13 | 25 | 28 | 0 | 3 | 0 |
| chr14 | 35 | 35 | 0 | 14 | 0 |
| chr15 | 26 | 34 | 2 | 22 | 19 |
| chr16 | 36 | 24 | 0 | 8 | 0 |
| chr17 | 28 | 22 | 0 | 10 | 7 |
| chr18 | 12 | 15 | 0 | 18 | 0 |
| chr19 | 6 | 5 | 0 | 1 | 7 |
| chr20 | 19 | 13 | 0 | 10 | 2 |
| chr21 | 12 | 3 | 0 | 4 | 0 |
| chr22 | 2 | 6 | 0 | 4 | 0 |
| chrX | 17 | 13 | 0 | 6 | 0 |

**Table 2:** Number of CCDs containing at least one solid linear minor in each chromosome, for each of the datasets. The percentage refers to the total number of CCDs on that chromosome in the given dataset. For GM12878 the (LR) and (IS) indicate the long-read ChIA-PET dataset or in situ ChIA-PET respectively.

|  | GM12878 |  | GM12878 |  | H1ESC |  | HFFC6 |  | WTC11 |  |
| --- | --- | --- | --- | --- | --- | --- | --- | --- | --- | --- |
| chr1 | 26 | 13.7% | 13 | 6.0% | 6 | 2.9% | 14 | 6.8% | 32 | 12.9% |
| chr2 | 43 | 24.0% | 20 | 11.2% | 1 | 1.7% | 20 | 13.0% | 29 | 13.4% |
| chr3 | 25 | 15.1% | 22 | 12.2% | 1 | 2.2% | 15 | 11.5% | 3 | 2.3% |
| chr4 | 33 | 27.7% | 11 | 8.9% | 0 | 0.0% | 10 | 13.3% | 0 | 0.0% |
| chr5 | 26 | 19.7% | 18 | 13.7% | 1 | 2.3% | 12 | 11.3% | 1 | 2.2% |
| chr6 | 28 | 20.4% | 8 | 5.7% | 0 | 0.0% | 12 | 9.8% | 2 | 4.3% |
| chr7 | 22 | 16.9% | 16 | 12.3% | 0 | 0.0% | 8 | 8.5% | 2 | 4.9% |
| chr8 | 29 | 26.1% | 16 | 15.5% | 0 | 0.0% | 12 | 14.3% | 1 | 6.2% |
| chr9 | 20 | 21.3% | 13 | 12.4% | 0 | 0.0% | 9 | 10.1% | 3 | 7.5% |
| chr10 | 10 | 9.1% | 16 | 12.9% | 0 | 0.0% | 19 | 18.4% | 3 | 4.7% |
| chr11 | 14 | 13.0% | 10 | 7.0% | 0 | 0.0% | 8 | 6.5% | 1 | 1.1% |
| chr12 | 21 | 18.1% | 20 | 16.4% | 0 | 0.0% | 15 | 15.3% | 3 | 4.7% |
| chr13 | 20 | 28.6% | 14 | 22.6% | 0 | 0.0% | 3 | 11.1% | 0 | 0.0% |
| chr14 | 11 | 15.3% | 7 | 9.5% | 0 | 0.0% | 7 | 12.7% | 0 | 0.0% |
| chr15 | 5 | 6.8% | 8 | 9.4% | 1 | 2.3% | 6 | 8.7% | 3 | 6.2% |
| chr16 | 11 | 16.2% | 11 | 14.3% | 0 | 0.0% | 6 | 8.7% | 0 | 0.0% |
| chr17 | 7 | 8.3% | 10 | 8.6% | 0 | 0.0% | 5 | 5.1% | 4 | 4.3% |
| chr18 | 7 | 12.7% | 4 | 6.5% | 0 | 0.0% | 4 | 13.3% | 0 | 0.0% |
| chr19 | 4 | 6.9% | 3 | 3.4% | 0 | 0.0% | 1 | 1.2% | 3 | 3.8% |
| chr20 | 6 | 12.8% | 8 | 11.9% | 0 | 0.0% | 5 | 8.1% | 2 | 6.2% |
| chr21 | 4 | 17.4% | 1 | 5.9% | 0 | 0.0% | 2 | 14.3% | 0 | 0.0% |
| chr22 | 2 | 6.1% | 5 | 10.4% | 0 | 0.0% | 4 | 10.8% | 0 | 0.0% |
| chrX | 8 | 8.0% | 3 | 6.0% | 0 | 0 | 2 | 15.4% | 0 | 0.0% |

### 3.6 Example of a linked structure

To give a more direct understanding of the actual minors and their relationship with the 3D structure we provide an example one solid linear K_6_ minor that was found in the in situ ChIA-PET for GM12878, on chromosome 10, between coordinates 86397456 to 86587768. A 3D model of the region is presented in Figure 2. This minor was selected because of the small number of edges in the graph (143 edges, 84 nodes), which makes the visualization more clear. Notice how the brown and green segments form a closed “loop” (i.e., a closed circuit, not to be confused with a chromatin loop), through which purple and red parts of the strand pass - these cannot be disentangled from one another, i.e., are linked. Note, that it is not possible to deduce from the graph which segments will form this link, as there are many geometric realizations of the graph - we only know that it must happen in this region.

**Figure 2:**
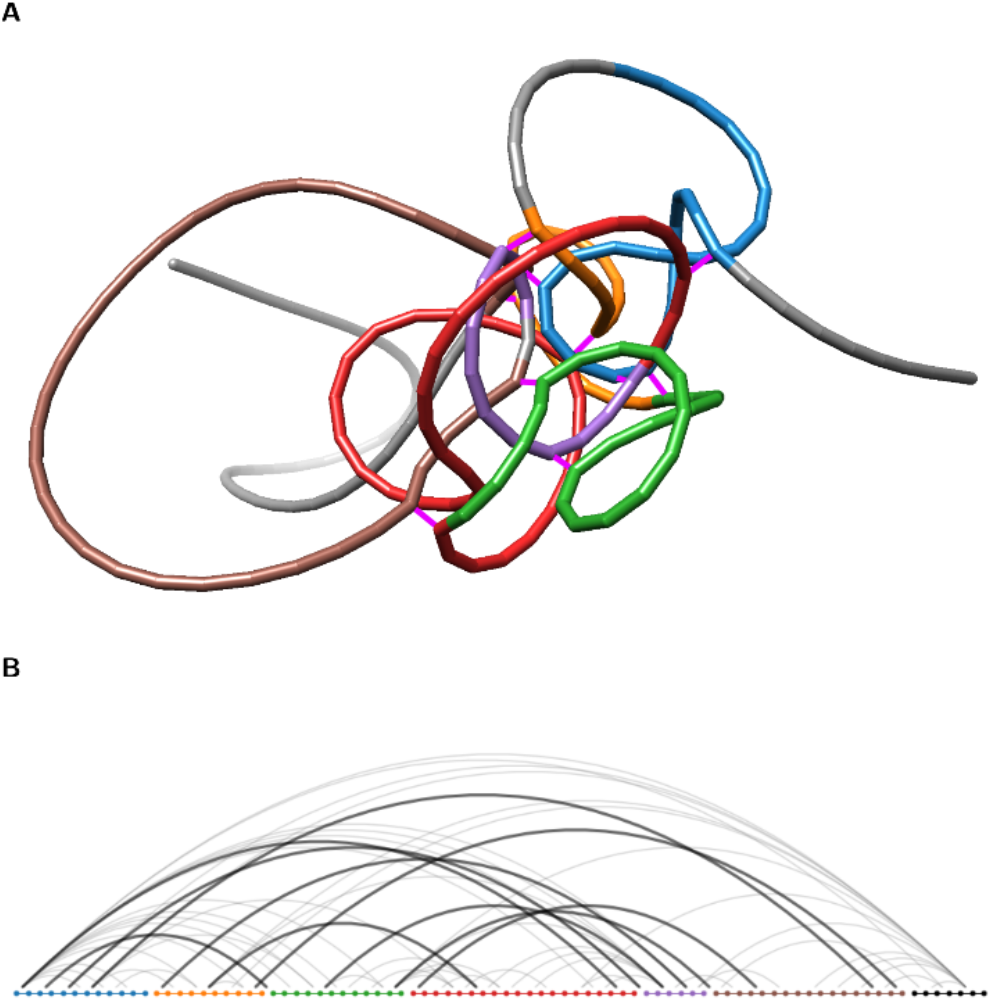
An example CCD, located at chr10:86397456-86587768, in which a linear minor was found. A) A 3D model of region created using Spring Model tool (Kadlof, Rozycka, & Plewczynski, 2020). The strand colors represent regions that collapse into a single vertex in the minor, while gray represents parts of chromatin not belonging to the minor. Fuchsia lines represent ChIA-PET interactions. The brown and green segments form a closed loop, through which purple and red parts of the strand pass - this is the linking in this particular geometric realization of the linked graph. B) Graph representation of the region. The dots represent the linear order of the vertices on the chromatin strand, colored as above. The arcs represent all ChIA-PET interactions in the region, with bold ones being the edges participating in the minor.

### 3.7 Topology of linked CCDs

We will now look at the graph-theoretic characterization of the regions containing the minors, focusing on GM12878. We compare the median values for each of the measures between the regions, in which at least one solid linear minor was found, and those without any minors. We will call the former “linked CCDs” and the others as “non-linked’.” To make the comparison reliable we excluded CCDs which were either very large or very small in terms of the size of the graph. First, we discarded the CCDs that have less than 6 nodes or less than 15 edges, as they could not contain a K_6_ minor. Moreover, we discarded CCDs above 10000 edges, for which the averages of centrality measures could be skewed (0.1% CCDs have more than 4750 edges). We perform the comparison for each dataset separately with the Mann-Whitney U test and use Bonferroni correction. For the results of all tests: test statistics, p-values and medians for all cells are reported in Table 3.

**Table 3:**
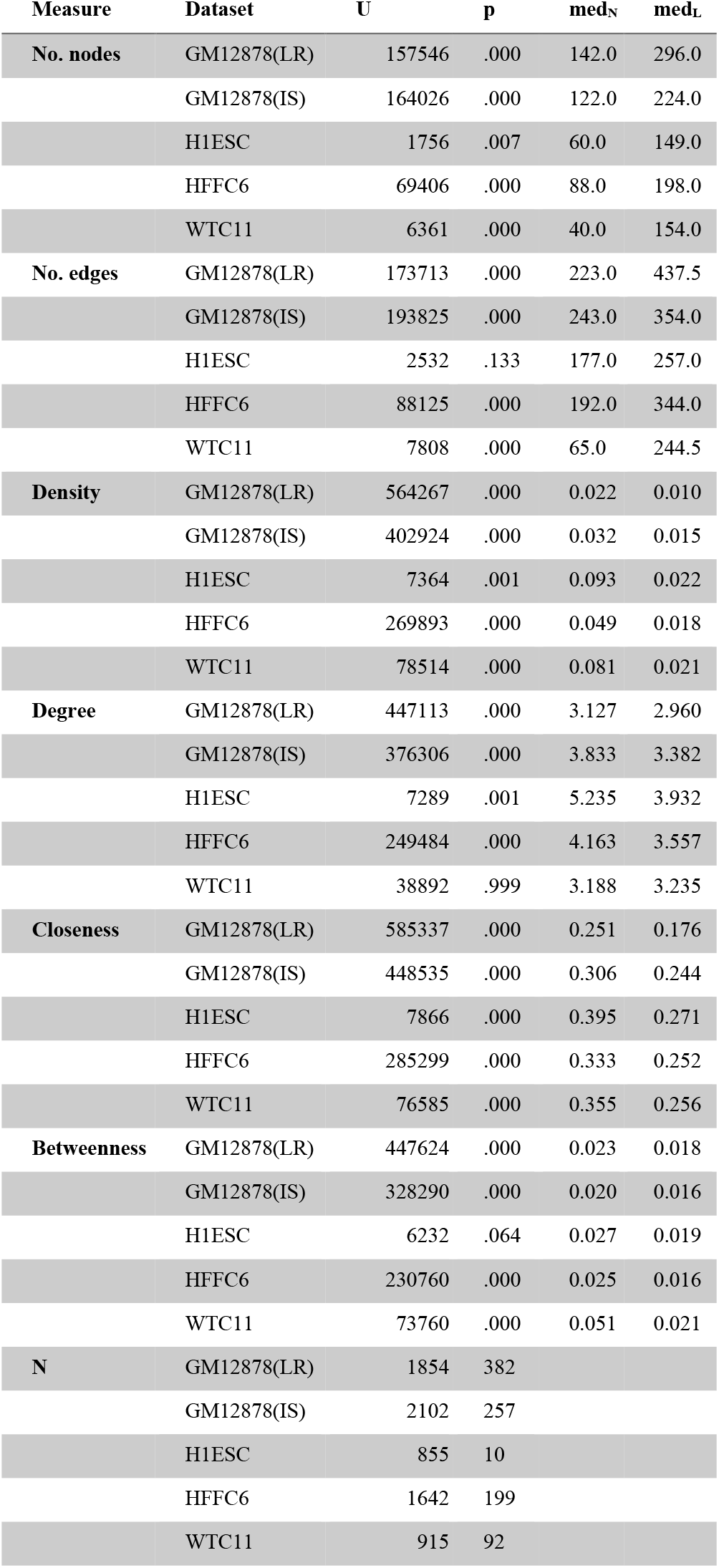
Results of each U Mann-Whitney tests performed to differentiate between linked and non-linked regions. Each graph measure is provided in separate set of rows. The columns are: U - the test statistic, p - the p-value, med_N_ - median value of a given measure for the CCDs without links, med_L_ - median value of for the linked CCDs. The group sizes in each sets are provide in the final set of rows. For GM12878 the (LR) and (IS) indicate the long-read ChIA-PET dataset or in situ ChIA-PET respectively.

The linked CCDs tend to be less dense, while having more nodes and edges in terms of absolute values. In case of GM12878, the medians of the numbers of nodes are 296 vs 140 (linked vs non-linked, p < .001) and 224 vs 122 (p < .001) for long-read and in situ ChIA-PET respectively. The densities, however, drop from 0.022 to 0.010 (p < .001) for long-read ChIA-PET and from 0.032 to 0.015 for in situ ChIA-PET (p < .001). This is interesting, given that denser graphs would seem to have more opportunity to randomly create clique structures.

Next, we find that both average closeness and average betweenness centrality in linked CCDs are lower than in non-linked. For long-read ChIA-PET the medians of closeness centrality are 0.25 for linked CCDs and 0.18 for non-linked (p < .001), and for in situ ChIA-PET their values are 0.31 for linked CCDs and 0.24 for non-linked CCDs (p < .001). The betweenness centrality medians are 0.023 vs 0.018 (p < .001) for long-read ChIA-PET, and 0.020 vs 0.016 (p = .020) or in situ ChIA-PET. The average degree also tends to slightly be lower in linked CCDs, in concordance with their lower density, being 3.13 vs 2.96 (p < .001) for long-read ChIA-PET and 3.83 vs 3.38 (p < .001) for in situ ChIA-PET.

The above comparisons hold true for four out of five datasets, except the H1ESC cell line, in which an exceedingly small number of minors found (only in 10 CCDs) (we provide the statistics nevertheless for the sake of completeness) - still, the tendencies in the medians are the same. Also, the degree did not vary significantly for the WTC11. The distributions of the graph-theoretic measures illustrating the comparisons are provided in Figure 3. These observations paint an overall characterization of the linked CCDs: they are regions with many redundant connections, as indicated by low betweenness; These connections do not, however, collapse the entire region, but only local tightly packed bundles exist - concordant with low closeness.

**Figure 3:**
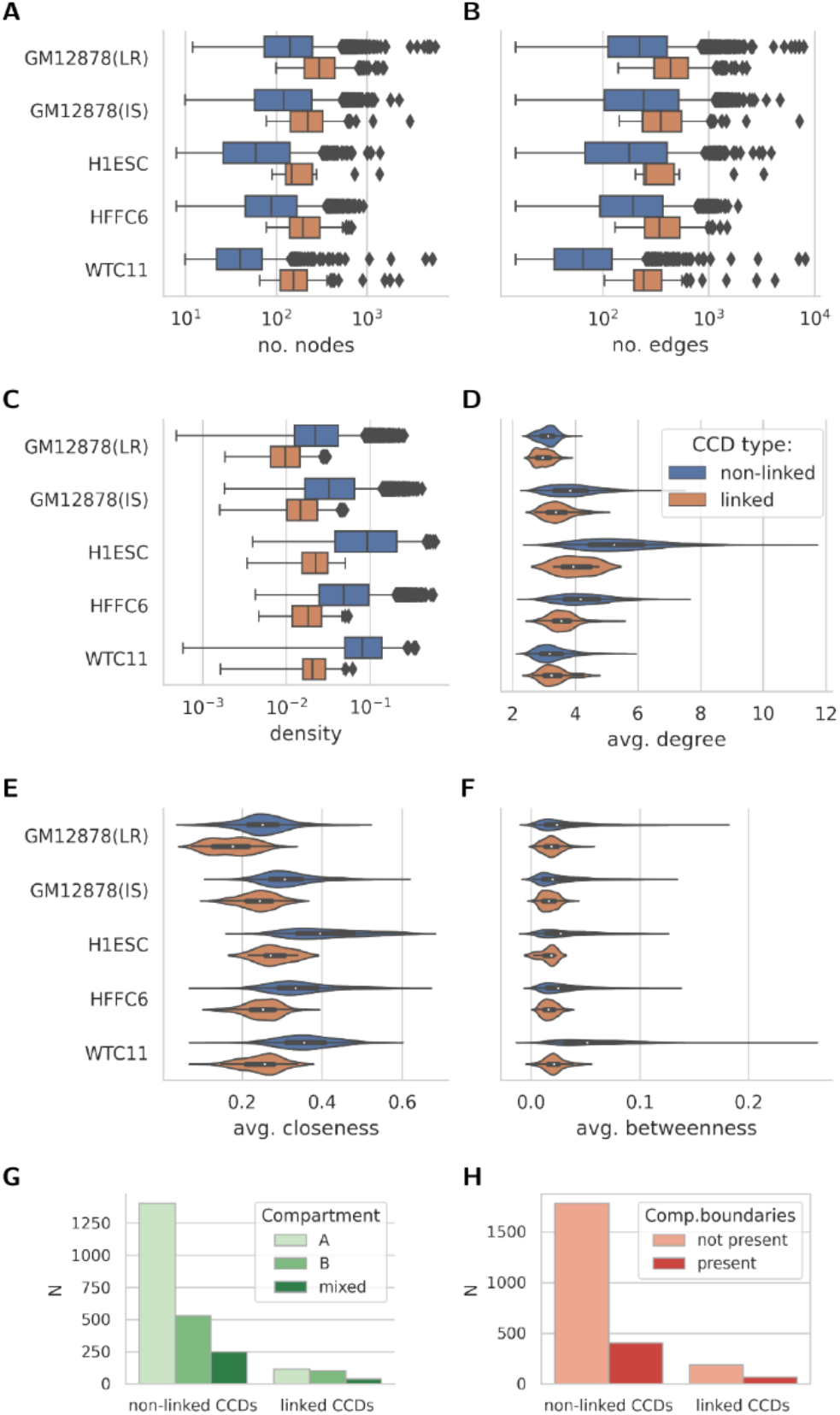
The distributions of graph properties for linked (orange) and non-linked (blue) CCDs, compared for each of the datasets separately. The first three box plots show A) the size of CCD graph in terms of number of vertices, B) number of edges, and C) density. In these plots the scale on the x axis is logarithmic. The next three violin plots show the per-node measures averaged within a CCD graph: D) average degree, E) average closeness centrality, F) average betweenness centrality. The last three plots show the counts of linked and non-linked CCDs by G) their assignment to chromatin compartments H) Presence of compartments boundaries within or near (< 50kb) them.

Finally, it is important to point out that the density in linked CCDs is lower than in non-linked, which indicates that the appearance of the minors is a deeper phenomenon, not explained simply by concentration of many interactions in a smaller region.

### 3.8 Linking and chromatin compartments

We obtained genomic coordinates of compartments discovered using in situ Hi-C experiments, for the GM12878 cell line, from 4DNucleome data portal (source lab: BCM) (Reiff, et al., 2021) (Dekker, et al., 2017). The resolution of the compartment data was 250kb, i.e., every segment of that length could have a compartment assigned. Our initial hypothesis was that the minors might aid in maintaining compartmentalization, so linked CCDs would be enriched in compartment boundaries, and would span multiple compartments more often.

To test this, for each region we calculated the proportion (expressed as a percentage) covered by either compartment. We then attributed each CCD to compartment A if the proportion of its length covered by compartment A was higher than 60%, to the B compartment if coverage by B was higher than 60% and marked any others as “mixed”. To complement this division, we calculated all points of change between compartments (1335 in total), and mapped them to CCDs with 50kb tolerance, dividing the CCDs into those either with or without such boundaries. The results show, that linked regions all almost equally to either compartment A or B, unlike non-linked regions - the chi-squared test for the contingency table yielded χ^2^(2) = 38.3, p < .001. The linked regions also contain slightly more compartment boundaries than expected (26% observed vs 19% expected, χ^2^(1) = 8.29, p = .005). Figure 3 shows the contingency tables for compartment assignment test and boundaries test, in the form of bar plots.

Overall, the results point towards the proposed enrichment, but not overwhelmingly.

### 4 Conclusion

We have described an algorithm capable of efficiently searching real-world genomic data for the presence of linked structures, which may play a role in spatial organization of chromatin. Our method found hundreds of regions with possible linking both for long-read and in situ ChIA-PET data in GM12878, and in three other human cell lines for the in situ method, with the numbers up to 1091, depending on the dataset. We found that it is rather structure than density of the region that predicts the presence of linking. The graph-theoretic characteristics of the linked regions distinguish them from other regions. An ideal clique has maximum possible closeness, as every node is connected to every other. For the same reason, the betweenness is minimal, as no route requires an intermediary. Low closeness in minor regions indicates that the groups of nodes are subdivided in such a way, that keeps relatively large number of nodes far apart, while retaining the redundancy of connections of the clique structure. These findings together support the view of structures that do not emerge simply from concentration of contacts in one place.

In the present study we searched only for linear minors i.e., minors with vertices occupying consecutive parts of chromosome. Our approach, however, provides another version of the algorithm: one searching for non-linear minors as well. While it could detect more linked structures, it has significantly larger computational complexity, making it unfeasible for larger datasets like the ones we used. Nevertheless, the linear minors alone provide enough data for statistical assessment of the differences between linked and non-linked regions, even though the number of minors would be higher.

One must bear in mind that the presence of the actual linked structure is contingent on the edges forming a minor being simultaneously present within a cell. Since our research is based on aggregated population data, it would be valuable to confirm results using a method that delivers single-cell data. However, the presently available methods cannot guarantee to capture all contacts, which severely limits the detection capability.

There are several implications of the presence of links, which need to be investigated in the future. Firstly, suppose that a linked structure is present in each chromosome. After breaking and reconnecting the DNA chain, e.g., using a topoisomerase, a link can be broken, but then it will reappear in a different place. This might be, potentially, a mechanism regulating gene expression in a cell. Secondly, mutations might destroy or create a linked structure, in a similar fashion that it can disrupt TAD boundaries (Valton & Dekker, 2016), potentially causing disease. On the other hand, a mutation that creates a linked structure might inhibit coding some important proteins or obstruct adaptation mechanisms. Finally, the presence of a rare contact or an absence of a commonly occurring contact might create or destroy some linked structure and lead to a different behavior of a cell, without affecting the DNA chain. To summarize, the proposed algorithm is a first step towards verification if intrinsic linking of chromatin is a viable mechanism for physically organizing the genome in the nucleus and regulating its function.

